# Similarity Measure for Sparse Time Course Data Based on Gaussian Processes

**DOI:** 10.1101/2021.03.03.433709

**Authors:** Zijing Liu, Mauricio Barahona

**Affiliations:** Department of Brain Sciences, Imperial College London, London, UK; Department of Mathematics, Imperial College London, London, UK

## Abstract

We propose a similarity measure for sparsely sampled time course data in the form of a loglikelihood ratio of Gaussian processes (GP). The proposed GP similarity is similar to a Bayes factor and provides enhanced robustness to noise in sparse time series, such as those found in various biological settings, e.g., gene transcriptomics. We show that the GP measure is equivalent to the Euclidean distance when the noise variance in the GP is negligible compared to the noise variance of the signal. Our numerical experiments on both synthetic and real data show improved performance of the GP similarity when used in conjunction with two distance-based clustering methods.

## 1 INTRODUCTION

Time course data are used widely for the empirical study of dynamical processes in many areas of research in the natural and social sciences [Keogh and Kasetty, 2003]. Traditionally, much research has been devoted to the characterisation of time series in relation to the originating dynamical process from different viewpoints, from the deterministic to the stochastic [Brillinger, 1981, Barahona and Poon, 1996, Chatfield, 2003].

More recently, time series have also been considered from the perspective of data science. One of the key questions in many applications is to assess the (dis-)similarity between time courses, with a view to perform time series classification or clustering [Liao, 2005, Son and Baek, 2008, Górecki, 2014, Peach et al., 2019, Fulcher and Jones, 2014]. A simple way to deal with finite, discretely-sampled time courses is to treat them as vectors [Yao et al., 2005, Hedeker and Gibbons, 2006], i.e., a time series sampled at *t* time points 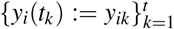 is described by a *t*-dimensional vector **y**_*i*_ with coordinates *y_ik_*. A simple dissimilarity measure between two time series **y**_*i*_ and **y**_*j*_ is then given by the Euclidean (*ℓ*^2^) distance:

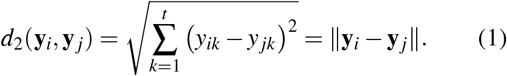

Alghouth the Euclidean distance is widely used due to its simplicity, in some applications, one may be more interested in the trend of how the data changes across time rather than the absolute differences. To capture this, a frequently used dissimilarity measure is based on Pearson’s correlation coefficient, 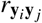:

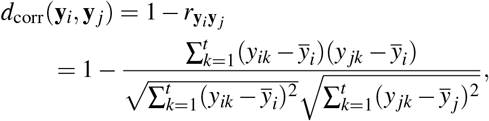

where 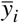 and 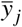 are the means of each time series. Note that these two common measures are point-to-point matching, and insensitive to the re-ordering of the time points and the spacing between the sampling times. Hence these measures cannot capture information associated with the time indices, which can be important in applications.

To remedy the limitations of point-wise measures, alternative measures of dissimilarity in time series have been proposed, including Dynamic Time Warping (DTW) [Keogh and Ratanamahatana, 2005] and the Edit Distance on Real Sequences (EDR) [Chen et al., 2005]. These methods can cope with uneven sampling and use information from the time indices, yet they can be algorithmically complex and are not well suited for applications with a large number of short, sparsely sampled time courses. Examples of this type of data are common in home price, marketing or ecommerce data in economics and finance [Fan et al., 2011], longitudinal electronic healthcare records in healthcare [Perotte and Hripcsak, 2013], genomics and proteomics data in life science [Ndukum et al., 2011, Kayano et al., 2016], and functional magnetic resonance imaging [Smith, 2012]. For instance, in cellular biology, ‘omics’ experiments measure the expression level of large numbers of genes, proteins or metabolites in cells over time. Such datasets contain tens of thousands of time courses (i.e., the number of genes or proteins) but the length of each time course is very short (5-15 time points) due to the high cost of experiments. Furthermore, the time samplings are usually uneven since experiments are designed to capture trends in cellular evolution and responses to stimuli. These constraints are typical of many biological experimental settings.

In this paper, we introduce a similarity measure for time course data based on Gaussian processes (GPs) [Rasmussen and Williams, 2006], which is applicable to sparse, inhomogeneously sampled, high-dimensional datasets. To retain information from the sampling times in the data, we model the time courses as continuous functions using GPs, and define a similarity measure in the form of a log-likelihood ratio between GP models. The GP similarity is computationally simple and suitable for high-dimensional datasets with a large number of short time courses. We also show that the GP similarity measure is equivalent to the Euclidean distance when the noise variance in the GP model is negligible compared to the signal variance. We apply the GP similarity measure as the basis for distance-based clustering methods in both synthetic and real time course data, and show improved robustness to measurement noise and to sampling inhomogeneity.

## 2 RELATED WORK

As a non-parametric model, GP is a flexible and efficient tool for time-dependent data modelling. Using the fact that a GP defines a reproducing kernel Hilbert space (RKHS), Lu et al. [2008] proposed a RKHS-based distance for time series defined as the Bregman divergence between the two posterior GPs. This distance has a closed form: it is the squared norm of the posterior mean functions in the RKHS induced by the GP. However, the Bregman divergence does not reflect the uncertainty of the data, since it only depends on the posterior mean function. Hence the Bregman divergence can perform poorly in the presence of noise in the data, as we show below.

GPs have been previously applied to time series of gene expression to detect differentially expressed genes [Stegle et al., 2010, Kalaitzis and Lawrence, 2011] and to infer the dynamics of transcriptional regulation [Lawrence et al., 2007, Gao et al., 2008]. In Kalaitzis and Lawrence [2011], the fitted GP model for each gene is compared to a noise model in order to rank the time courses and find differentially expressed genes. Here, we use the construction of GPs differently, and show that the likelihood ratio between two GP models provides a robust similarity measure for time courses. In the next section, we will introduce the GP model for time course data.

## 3 GAUSSIAN PROCESS MODEL FOR TIME COURSE DATA

A Gaussian process is a collection of random variables over the index set 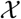 such that any finite collection of the random variables follows a multivariate normal distribution [Rasmussen and Williams, 2006]. Therefore, a Gaussian process, denoted 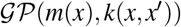, is characterised by the mean function *m*(*x*) and the covariance function *k*(*x,x*′), where 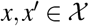. Here we will consider time-dependent variables; hence the index set is the positive real line describing time: 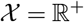.

We will model the underlying true signal as a Gaussian process over time:

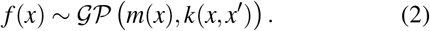

For simplicity, we take the mean function to be zero (*m*(*x*) = 0) and we use the squared exponential covariance function

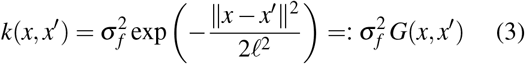

where *ℓ* is a characteristic length-scale, 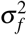 is the signal variance and we use *G*(*x, x*′) to denote the Gaussian kernel. The observations of the time dependent variable are then noisy samples of the GP:

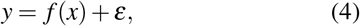

where the additive noise *ε* is Gaussian with zero mean and variance 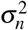.

Let us consider a time-dependent variable *y* given by (4) sampled at *t* time points *X* = [*x*_1_, …, *x_t_*]^*T*^ and let us compile the observations into a *t*-dimensional vector **y** = [*y*_1_, …, *y_t_*] ^*T*^. Under our assumptions, the covariance function of the noisy observations *y* is given by:

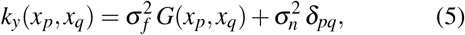

where *δ_pq_* is the Kronecker delta. Equivalently, the *t* × *t* covariance matrix of the observations **y** is:

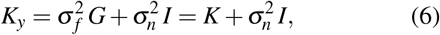

where *G* is the Gaussian kernel matrix with elements *G_pq_* = *G*(*x_p_, x_q_*), *I* is the identity matrix of dimension *t*, and *K* is the covariance matrix for the noiseless samples with elements *K_pq_* = *k*(*x_p_, x_q_*).

The three hyperparameters of the Gaussian process are therefore ***θ*** = (*ℓ, **σ**_f_, **σ**_n_*), and can be learnt from the data (*X*, **y**) by maximising

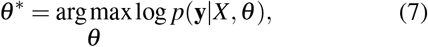

where, in this case, the log-marginal likelihood has the explicit form:

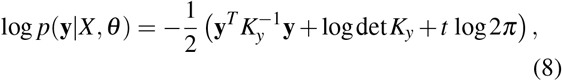

with det *K_y_* denoting the determinant of *K_y_*. This expression can be maximised using gradient-based methods [Rasmussen and Williams, 2006].

## 4 SIMILARITY MEASURE FOR TIME COURSE DATA BASED ON GAUSSIAN PROCESS

Let us consider a sparse time course dataset consisting *N* short time courses sampled at *t* time points: **y**_*i*_ ∈ ℝ^*t*^, *i* = 1,…, *N*. The dataset is referred to as sparse, due to the fact that the number of time courses is much larger than the length of each time course (*N* ≫ *t*). Although the GP model does not require all the time series to be measured synchronously, for simplicity, we first introduce the similarity measure for the case where all time courses are sampled at the same time points *X* = [*x*_1_, …, *x_t_*]^*T*^. We discuss the asynchronous case later.

Each of the time courses **y**_*i*_ is assumed to correspond to a noisy observation of a Gaussian process (4) with the same hyperparameters *θ*, which can be inferred by maximising the sum of the log marginal likelihoods of the time courses [Rasmussen and Williams, 2006]:

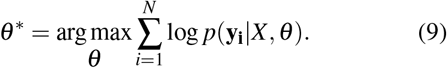

By inferring the hyperparameters *θ*^∗^, we obtain a nonparametric probabilistic model for the observed time courses, which we can use to define a GP-based similarity measure, as follows.

### 4.1 LIKELIHOOD RATIO AS A SIMPLE GP SIMILARITY MEASURE

Using the fact that the GP is a distribution of continuous functions over time, a similarity measure between two observed time courses **y**_*i*_ and **y**_*j*_ can be obtained by comparing two different possibilities as to how **y**_*i*_ and **y**_*j*_ could have been generated.

The first possibility is that the two time samples **y**_*i*_ and **y**_*j*_ are observations from *two different* functions sampled from the GP. In this case, the joint likelihood of **y**_*i*_ and **y**_*j*_ is just the product of the likelihoods of two time courses:

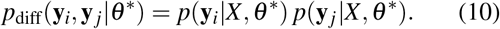

Using (8), it is easy to see that the log-likelihood can be rewritten as:

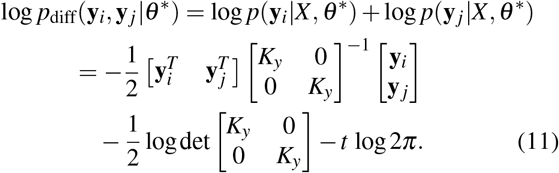

The second possibility is that **y**_*i*_ and **y**_*j*_ are observations from *the same* function sampled from the GP. In this case, the joint likelihood of **y**_*i*_ and **y**_*j*_ can be computed by considering the two time courses {*X*, **y**_*i*_} and {*X*, **y**_*j*_} to be replicate samples of one function:

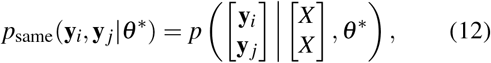

and the log-likelihood is then given by:

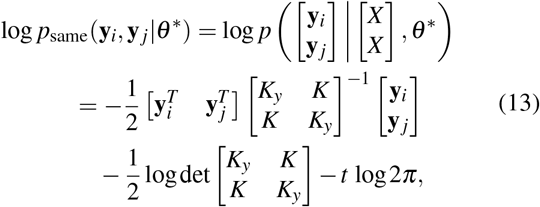

where we have used the fact that the additive noise *ε* in (4) is uncorrelated between the two time courses.

The likelihood (12) will be high if the two time courses are similar to each other (as in Fig. 1a), and will be small if the two time courses have different profiles (as in Fig. 1b). Hence for time courses with different profiles, the likelihood (10) explains better the data. The log of the ratio of the two likelihoods (12)–(10) (i.e., the difference between the log-likelihoods) is thus an indicator of the level of similarity between two time courses. This leads to our definition of the *GP similarity measure* between **y**_*i*_ and **y**_*j*_ as:

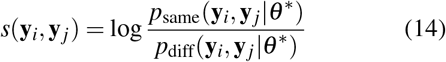

and it follows that:

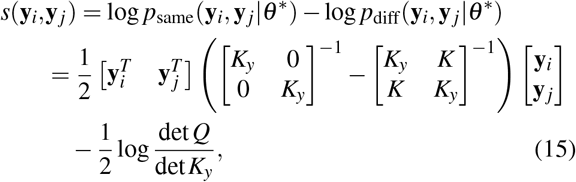

where *Q* is the Schur complement:

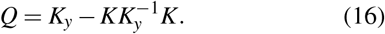

**Figure 1:**
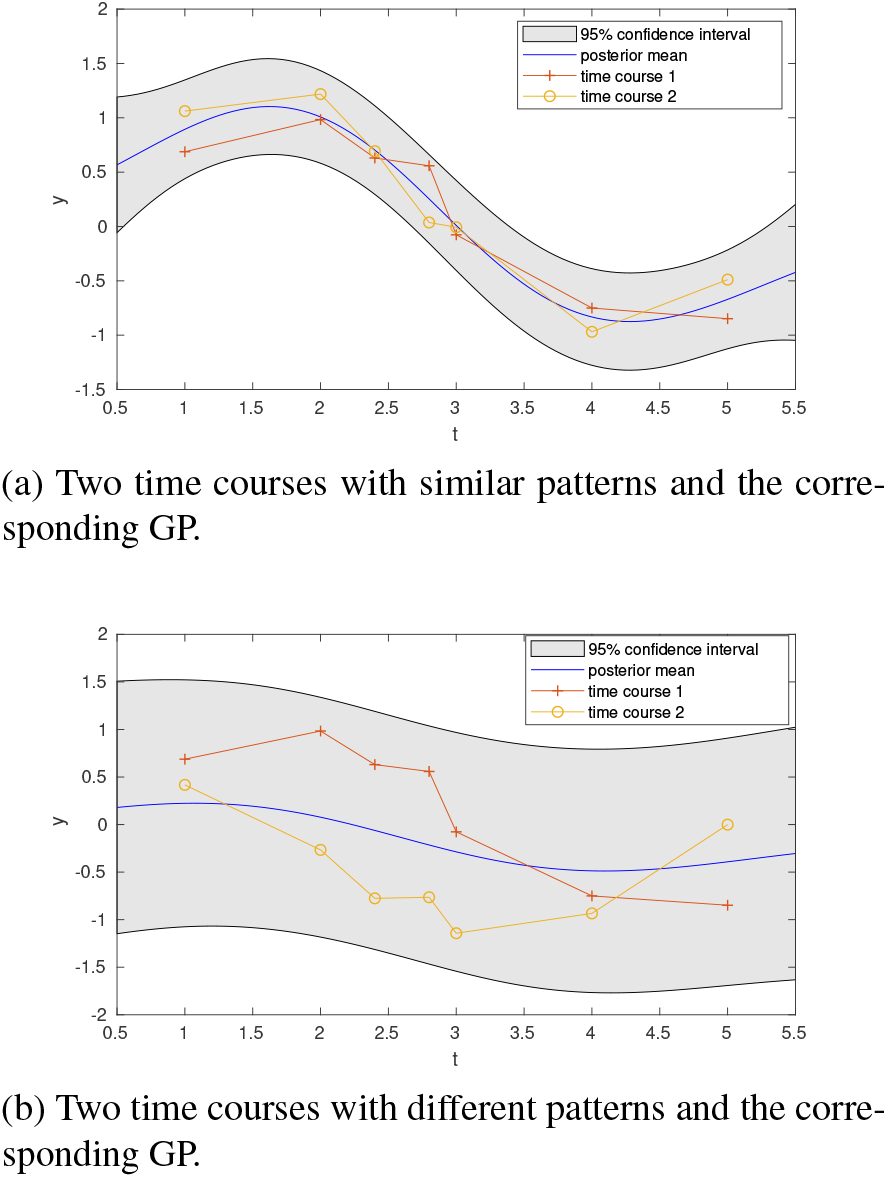
Two examples with two time courses that have: similar and (b) dissimilar profiles. In each case, we show the time courses and the mean and confidence interval of the Gaussian process (12) obtained according to the likelihood in (12).

Note that the likelihood ratio (14) has the same form as the Bayes factor [Kass and Raftery, 1995] in Bayesian model selection. From this perspective, our measure can be understood as the comparison between two models in two ways: on one hand, *s*(**y**_*i*_, **y**_*j*_) compares modelling two time courses with one single function of the GP versus modelling them with two independent functions of the GP; alternatively, it is easy to see that our measure (14) can be rewritten as

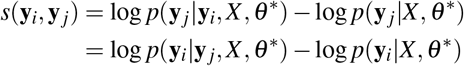

which is the difference between the log-likelihood of **y**_*j*_ based on the posterior GP given **y**_*i*_ compared to the prior GP without **y**_*i*_ being given. Hence the measure quantifies the improvement in the prediction of **y**_*j*_ that can be drawn by knowing **y**_*i*_. The measure is symmetric, so the same applies by exchanging **y**_*j*_ for **y**_*i*_.

### 4.2 EUCLIDEAN DISTANCE AS A LIMIT OF THE GP LOG-LIKELIHOOD RATIO

It can be shown that the Euclidean distance (1) stems naturally from the GP similarity (14) in the limit when the noise variance 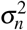 is much smaller than the signal variance 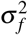.

To see this, recall the Neumann series [Stewart, 1998].

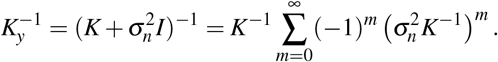

Noting that the Gaussian kernel matrix *G* is positive definite, we have that

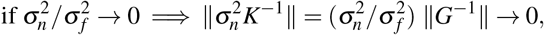

For small 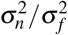, we thus take the first two terms of the expansion to 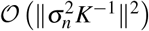:

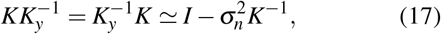

and the Schur complement (16) is approximated as:

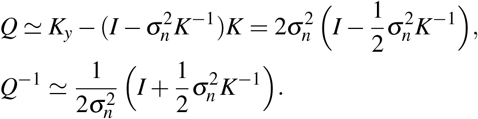

To approximate the GP similarity measure (15), we use block matrix inversion:

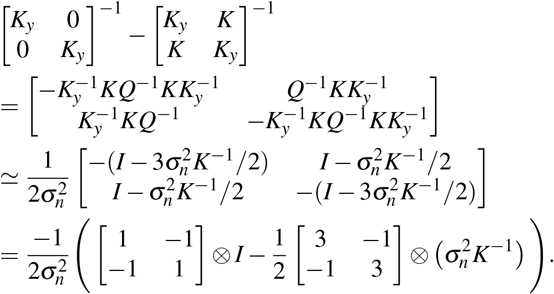

and we approximate the determinant:

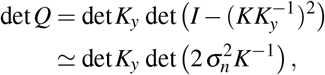

whence we obtain:

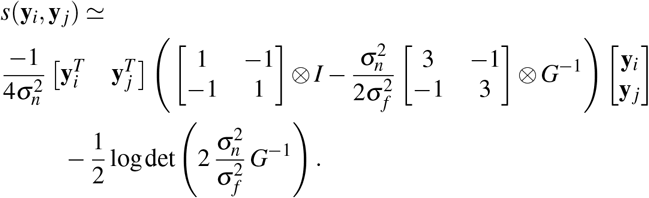

Here ⊗ denotes the Kronecker product and *G* is the Gaussian kernel matrix in (6). The approximation of the GP similarity measure (15) to first order is then:

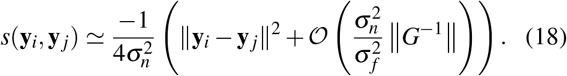

Therefore, if 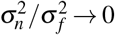 (i.e., when the variance of the noise is much smaller than the signal variance), the dissimilarity measure −*s*(**y**_*i*_, **y**_*j*_) is equivalent to the Euclidean distance ∥**y**_*i*_ − **y**_*j*_∥^2^. Note that this relationship holds not only for the Gaussian kernel *G*(*x, x*′) but for any positive definite kernel.

### 4.3 ASYNCHRONOUS TIME COURSES

Although, for simplicity of exposition, we have concentrated on the case of synchronous time sampling, the GP similarity measure is equally applicable to non-synchronous samples. In our derivations above, synchronous time points are only necessary to obtain the formal limit to the Euclidean distance in Eq. 18.

To see the applicability to non-synchronous samples, note that the likelihoods in Eq. 13 and Eq. 11 do not require the time courses to have the same time points. We can therefore consider *N* time courses denoted by 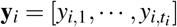 of length *t_i_*, sampled at (potentially) distinct time points 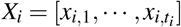 (*i* = 1,…,*N*). In this case, we can similarly model the time courses with GP and learn the hyperparameters by maximizing

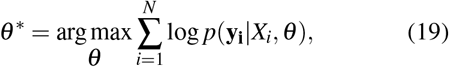

and it is then easy to write the two log-likelihoods:

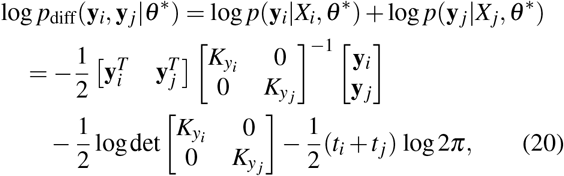

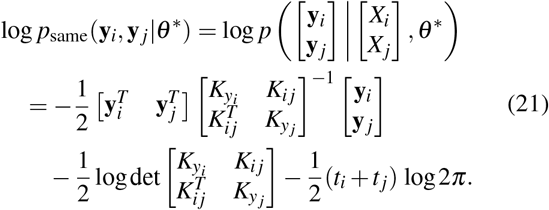

where 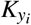 and 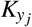 are the covariance matrices of **y**_*i*_ and **y**_*j*_ with sizes *t_i_* × *t_i_* and *t _j_* × *t _j_*, respectively; and *K_i j_* is the cross-covariance matrix between **y**_*i*_ and **y**_*j*_ with size *t_i_× t_j_*. The GP similarity between the two time courses can again be computed as the difference between the two log-likelihoods (20) and (21).

### 4.4 COMPUTATIONAL COMPLEXITY

In terms of computational complexity, fitting a GP model with *N* time courses of length *t* takes 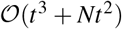 time. Computing pairwise similarities takes 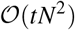 time. Since we deal with high-dimensional short time courses (*N* ≫ *t*), the total time for GP similarity would be approximately 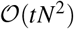, which is the same as for the Euclidean distance.

Extra computational time is needed for the asynchronised time courses, where all the time courses have a different covariance matrix. It results in a computational time of 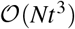 for model fitting and 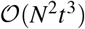 for computing the pairwise GP similarity if all the time courses are of average length *t*. This might be a limitation to applications with large *N* and *t*.

## 5 NUMERICAL EXPERIMENTS

To test the applicability of the proposed GP similarity measure, we have run numerical experiments on synthetic and real sparse time course data. Our results show that the similarity *s*(**y**_*i*_, **y**_*j*_) is more robust to observational noise than the Euclidean distance when used as the basis to cluster time courses using two standard clustering algorithms (hierarchical and spectral clustering). We also compare the performance of the GP similarity against the Bregman divergence and Dynamic Time Warping.

### 5.1 SYNTHETIC DATA

Our synthetic dataset is obtained by sampling from the three different time profiles shown in Fig. 2 with additive Gaussian noise. From each of the three time profiles, we generate 50 evenly-sampled time courses of length *t* = 15 with a given level of sampling noise. Our task is to cluster the 150 time series into 3 groups in an unsupervised manner. To do this, we compute the pairwise similarities (or distances) between the 150 time courses using the GP similarity (14). From the computed similarity matrix, we then cluster the samples using two well-known methods: (i) spectral clustering with *k*NN graph (*k* = 7) [Yu and Shi, 2003]; (ii) agglomerative hierarchical clustering with average linkage [Rokach and Maimon, 2005]. We then repeat the numerical experiment 100 times in each case. To evaluate the clustering performance, we use the normalised mutual information (NMI) [Vinh et al., 2010] against the known ground truth (i.e., the three profiles used to generate the data).

**Figure 2:**
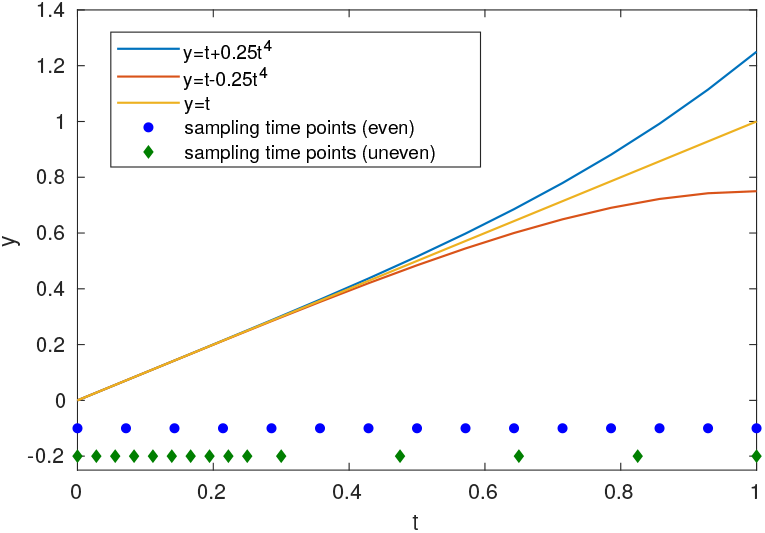
The three functions and the sampling time positions used to generate the synthetic data. The results of even sampling are in Fig. 3. The results of uneven sampling are in Fig. 4.

We then repeat the same procedure with three popular measures: the Euclidean distance, the Dynamic Time Warping (DTW) distance and the Bregman divergence in the RKHS. Given that the only varying ingredient is the similarity measure, the clustering performance reflects the quality of the similarity measure for this purpose.

Figure 3 shows the clustering quality (0 ≤ NMI ≤ 1) for evenly sampled time series with increasing levels of sampling noise achieved with both clustering methods and all four distance/similarity functions. As expected from (18), both Euclidean and GP similarity give comparable results for small sampling noise when using spectral clustering. Note, however, that even for small noise, the performance of the GP similarity is superior to the Euclidean distance when using hierarchical clustering (Fig. 3a), a method that is very sensitive to noisy data. Also as expected the DTW performs worse than the Euclidean distance for the synchronous case, since there is no need for alignment. The Bregman divergence does not perform well in this case because it measures the distance between the two continuous curves in the RKHS fitted from the time courses data and does not consider the uncertainty due to noise.

**Figure 3:**
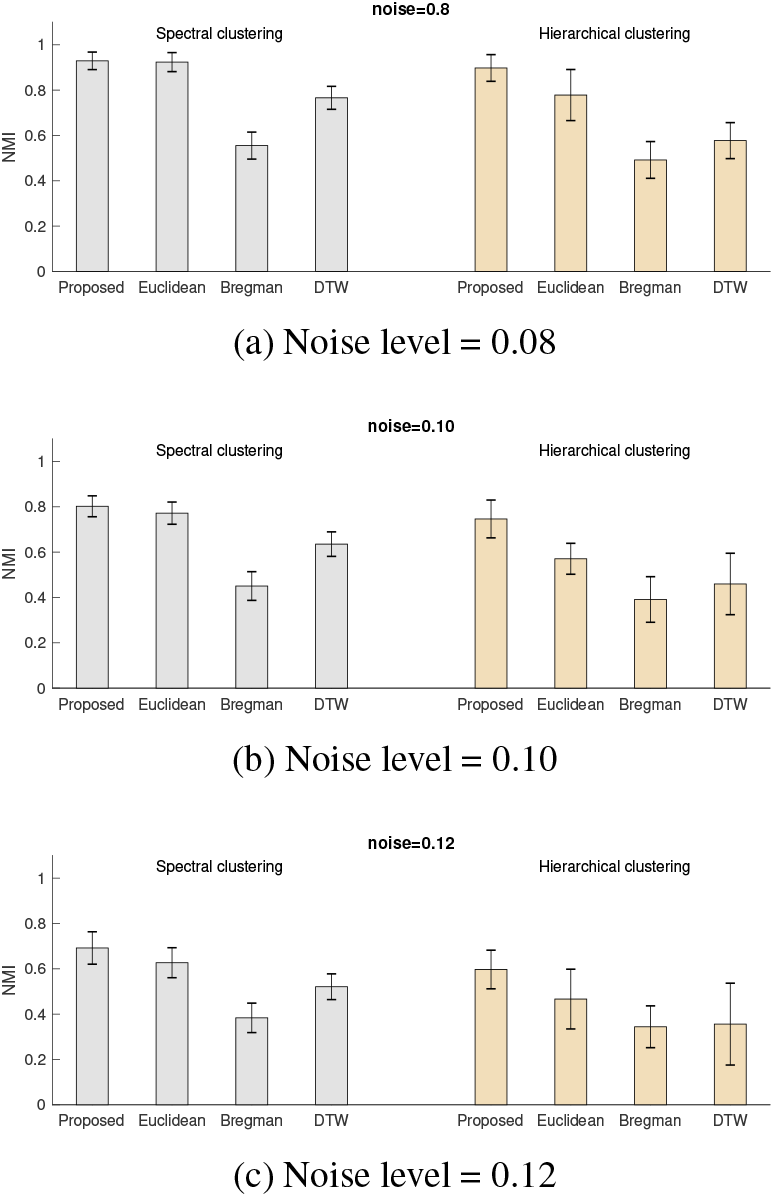
Clustering performance (measured as NMI) for spectral clustering with *k*NN and hierarchical clustering based on the proposed GP similarity (14), Euclidean distance (1), DTW and Bregman divergence in the RKHS with increasing noise levels from (a)-(c). The sampling time points are equally spaced between 0 and 1 (see Fig. 2).

As the observation noise increases, our GP similarity measure gains further advantage over the Euclidean distance for both clustering methods (Figs. 3b–3c). The p-values for a Wilcoxon rank-sum test between the NMI values of Euclidean and GP similarity with spectral clustering for the noise levels 0.08, 0.10 and 0.12 are 0.043, 2.5e-7 and 7.6e-16, respectively. The p-values for a Wilcoxon rank-sum test between the NMI values of Euclidean and GP similarity with hierarchical clustering for the noise levels 0.08, 0.10 and 0.12 are 1.4e-15, 1.7e-25, and 1.1e-20, respectively. Hence the GP similarity measure performs significantly better than the Euclidean distance for clustering with both algorithms.

In general, spectral clustering always performs better than hierarchical clustering, which is more sensitive to noise, and the best performance is obtained consistently using spectral clustering with GP similarity. Note that the performance obtained with spectral clustering using other metrics can be achieved at a lower computational cost using hierarchical clustering with GP similarity.

We also analyse a second set of 150 time series of length *t* = 15 collected from the same three functions in Fig. 2 but sampled inhomogeneously in time. The computational procedure is identical to the case of evenly sampled series described above. Figure 4 shows that the advantage of the GP similarity measure against the other distances is more prominent when the time points are unevenly sampled. This result highlights the fact that point-to-point similarities (such as the Euclidean distance) can miss important information contained in the long-term trends of the time profiles if the sampling is concentrated irregularly in particular time periods.

**Figure 4:**
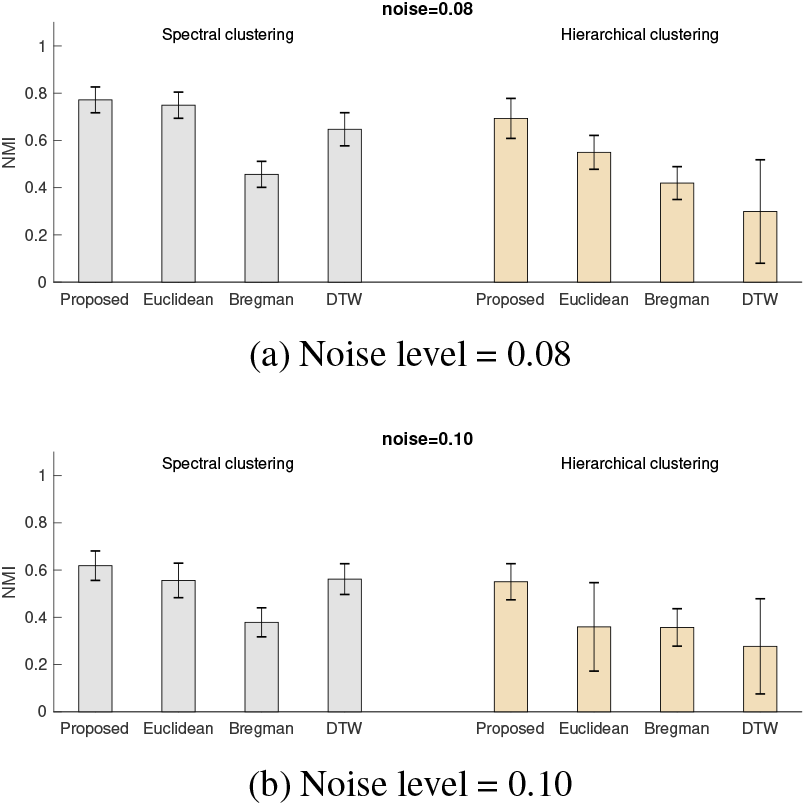
Clustering performance (measured as NMI) for spectral clustering with *k*NN and hierarchical clustering based on the proposed GP similarity (14), Euclidean distance (1), DTW and Bregman divergence in the RKHS with increasing noise levels from (a)-(b). In this case, the time points are irregularly sampled between 0 and 1 (see Fig. 2). The GP similarity consistently outperforms the other distances, especially for larger amounts of noise.

We next tested the non-synchronous case. We again generate a set of 150 time series of length *t* = 15 from the same three functions in Fig. 2. We then make the asynchronous time courses by randomly removing 6,7 or 8 time points from our 15-point time course data (Fig. 5a). In this case, the Euclidean distance is not defined. The proposed GP similarity achieves substantially better results than the Bregman divergence and DTW on these examples.

**Figure 5:**
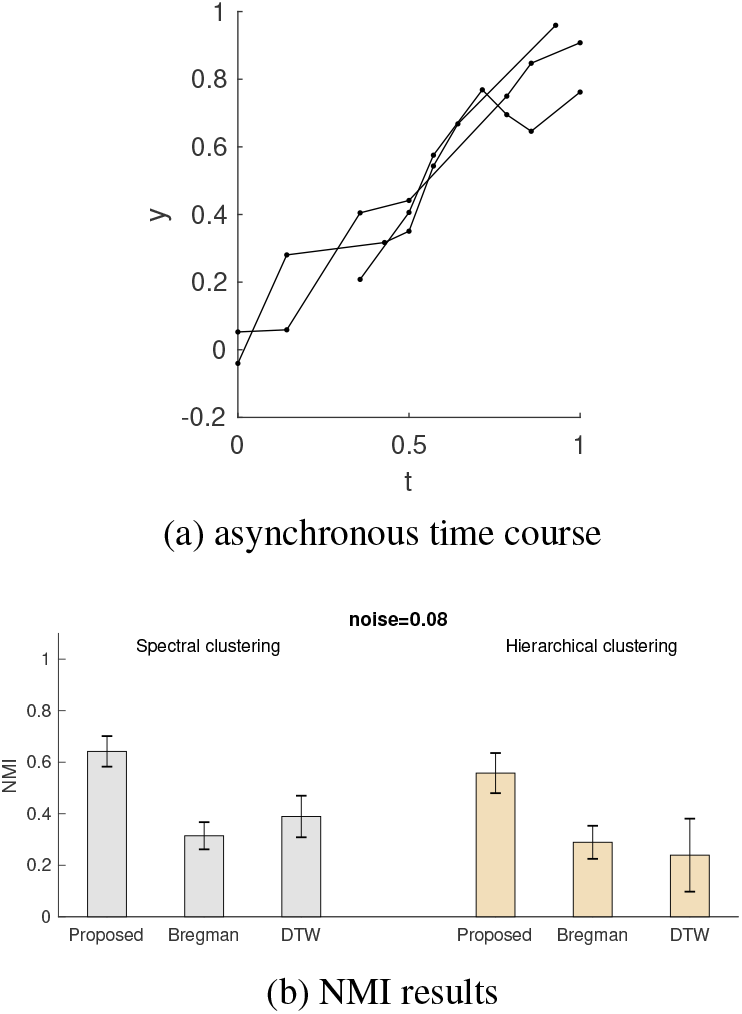
(a) Three sample time courses with asyncronised measurements with Noise level = 0.08. (b) Clustering performance (measured as NMI) for spectral clustering with *k*NN and hierarchical clustering based on the proposed GP similarity, DTW and Bregman divergence in the RKHS. The GP similarity consistently outperforms the other distances for both clustering methods.

In summary, our numerical experiments on synthetic data indicate improved performance of the GP similarity measure (14). As the observational noise decreases, the performance of the GP similarity measure is equivalent to the Euclidean distance (18). Although the GP similarity shows enhanced performance for both clustering methods, the improvement is larger for hierarchical clustering, in the presence of large amounts of noise, and under uneven time sampling. As a side comment, we also note that the inclusion of the noise variance 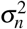 in the covariance function *K_y_* makes the GP similarity measure more robust for numerical computations, as it reduces numerical instability.

### 5.2 APPLICATION TO GENE EXPRESSION TIME COURSE DATA

We next tested the GP similarity measure on a real time course dataset which characterises the gene expressions during the process of cellular reprogramming where differentiated cells are reverted to stem cells [Di Stefano et al., 2014b]. In this dataset, gene expression values were measured at five time points: 0, 2, 4, 6 and 8 days after the cellular reprogramming started. Two biological replicates were measured at each time point. The full measurements include around 40,000 genes. Following standard practice, we select 1912 genes that are highly variable over time. We then follow the same procedure as for the synthetic data above to cluster the 1912 gene expression time courses to extract groups of genes that have similar time profiles during cellular reprogramming.

For this dataset, there is no known ‘ground truth’ against which to compare the obtained clusters. To assess the quality of the clustering, we use a biological score: a modified version of the biological homogeneity index (BHI). The BHI measures within-cluster homogeneity of the genes in terms of biological knowledge and is given by:

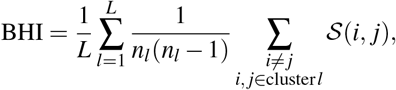

where *L* is the number of clusters, *n_l_* is the size of cluster *l*, and 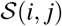 is a *biological similarity* between gene *i* and gene *j*. In the original BHI [Datta and Datta, 2006], the similarity between genes is either 0 or 1, which is an indicator function of two genes sharing any Gene Ontology (GO) terms. In order to better capture the biological information, we use an information-theoretic semantic gene similarity based on Gene Ontologies [Resnik, 1999, Lord et al., 2003, Yu et al., 2010]. For a given clustering, we compute the BHI of 1000 random clusterings with the same number and size of clusters. We then use the z-score of the BHI against the random clusterings as a level of significance for the obtained clustering.

Since the number of clusters is unknown, we compute the z-score of the BHI for clusterings with different numbers of clusters (Table 1). Again, we find that spectral clustering with GP similarity achieves the best performance. On the other extreme, hierarchical clustering based on Euclidean distances performs no better than random clustering, an indication that the gene expression data has high levels of noise *σ_n_*. As was the case for the synthetic data above, using GP similarity improves the performance of hierarchical clustering substantially, almost to a comparable level to the spectral method (but below). These results underscore the ability of the GP similarity to deal with noisy data.

**Table 1:** BHI z-scores of the two clustering methods with different number of clusters obtained by analysing the time courses of 1912 highly variable genes in the stem cell transcriptomic dataset of Refs. [Di Stefano et al., 2014a,b].

| Method | Similarity | No. of clusters |  |  |  |  |
| --- | --- | --- | --- | --- | --- | --- |
|  |  | 9 | 11 | 13 | 15 | 16 |
| Spectral | Proposed | 2.505 | **4.250** | 1.360 | 3.368 | 3.674 |
|  | Euclidean | 1.368 | 2.067 | 2.460 | 2.351 | 2.919 |
|  | DTW | 2.636 | 2.087 | 2.774 | 2.720 | 3.182 |
|  | Bregman | 4.229 | 2.725 | 2.652 | 3.617 | 3.520 |
| Hierarchical | Proposed | 1.182 | 2.384 | 2.366 | 1.849 | **3.353** |
|  | Euclidean | -0.090 | 0.074 | -0.266 | -0.215 | -0.319 |
|  | DTW | 0.542 | 0.566 | 0.499 | 0.735 | 0.752 |
|  | Bregman | 1.392 | 1.303 | 1.139 | 1.388 | 1.508 |

To illustrate visually the behaviour of the GP similarity, Figure 6 shows the time courses of three genes where the GP similarity and Euclidean distance behave differently. In our chosen examples, the time course of the *Samd14* gene (Fig. 6a) has the same Euclidean distance to both the *Fbin5* gene (Fig. 6b) and the *Serpine1* gene (Fig. 6c). On the other hand, according to our GP similarity, the *Serpine1* time course is more similar to *Samd14* than to *Fbin5*, in accordance with our visual expectation from the observed time course profiles. The codes and data are available online in https://github.com/barahona-research-group/BayesFactorSimilarity.

**Figure 6:**
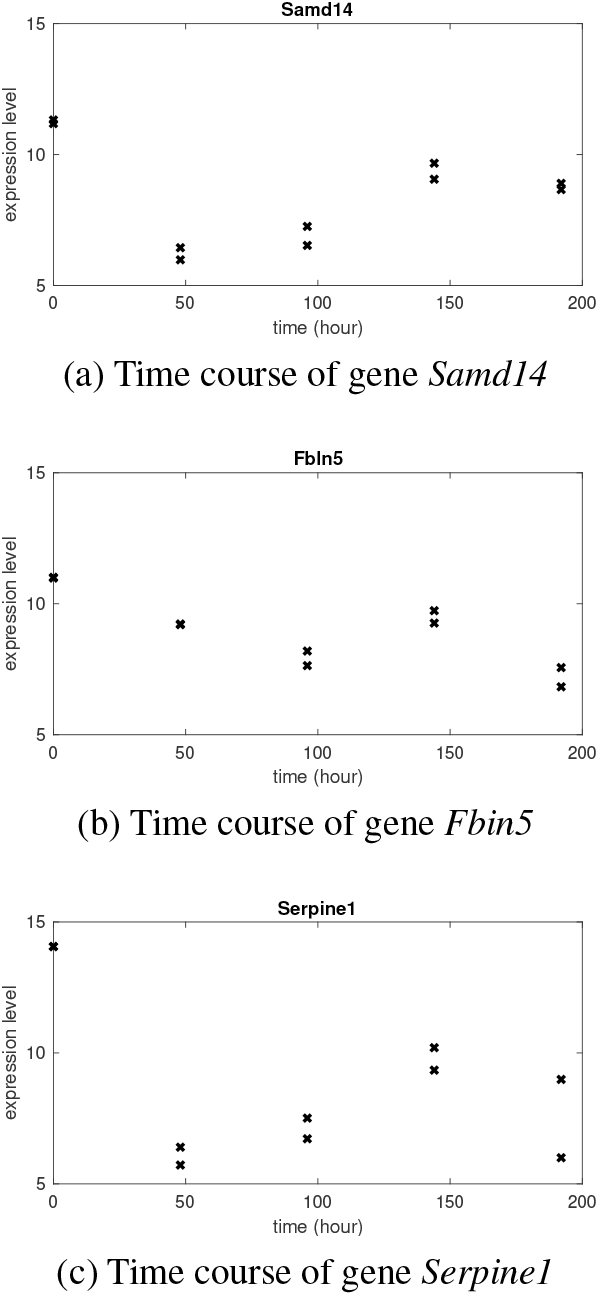
The time courses of three genes (each time course was measured on two bioogical replicates). The Euclidean distances from *Samd14* to *Fbin5* and *Serpine1* are both 5.09. In contrast, the GP dissimilarity (−*s*(**y**_*i*_, **y**_*j*_)) between *Samd14* and *Fbin5* (37.48) is larger than between *Samd14* and *Serpine1* (25.57), capturing the fact that the time profile of *Serpine1* is more similar to *Samd14* than to *Fbin5*.

## 6 CONCLUSION

In this paper, we have proposed a similarity measure for sparse time course data based on Gaussian processes. Modelling the time courses with a GP, we use the difference between two log-likelihoods (in the form of a Bayes factor) as a GP similarity measure. We show that the proposed measure is equivalent to the Euclidean distance in the limit where the noise variance in the observations is negligible compared to the signal variance. The proposed measure is computationally simple and can be easily extended to the cases when the time courses are not synchronously observed at the same time points.

Our numerical experiments show that the proposed measure has improved robustness to noise when used for data clustering with different clustering methods. The advantage of the proposed measure over the Euclidean distance is more noticeable with hierarchical clustering, under high noise, and with uneven time sampling.

We note two limitations of the proposed measure. First, the GP similarity is not a metric. Indeed, *s*(**y**_*i*_, **y**_*i*_) is not zero but instead, it gives an estimate of the level of noise in **y**_*i*_. Second, the high computational cost for the non-synchronous data may be a limitation if both the length *t* and number of time courses *N* are large. Sparse GP models can be used to overcome this limitation [Liu et al., 2020]. Although in this study we have only considered a GP over time as a one-dimensional space, future studies could generalise the GP similarity measure to data that can be modelled by GPs in a latent space, or on any other smooth manifold, further enhancing its applicability.

## Acknowledgements

This work was supported by the European Commission [European Union 7th Framework Programme for research, technological development and demonstration under grant agreement no. 607466]; and the Engineering and Physical Sciences Research Council [under grant EP/N014529/1 to M.B. funding the EPSRC Centre for Mathematics of Precision Healthcare].

